# Spatiotemporal dimensions of a reproductive life history trait in a spiny lizard radiation (Squamata: Phrynosomatidae)

**DOI:** 10.1101/2020.06.17.157891

**Authors:** Julián A. Velasco, Gustavo Campillo-García, Jesús Pinto-Ledezma, Oscar Flores-Villela

**Affiliations:** Centro de Ciencias de la Atmósfera, Universidad Nacional Autónoma de México. Mexico City, Mexico; Museo de Zoología “Alfonso L. Herrera”, Facultad de Ciencias, Universidad Nacional Autónoma de México. Mexico City, Mexico; Department of Ecology, Evolution, and Behavior, University of Minnesota, 1479 Gortner Avenue, Saint Paul, Minnesota 55108

**Keywords:** macroecology, macroevolution, life history, *Sceloporus*, cold-climate hypothesis

## Abstract

The ecological and evolutionary factors underlying life history trait variation is one of the most interesting topics in biology. Although many studies have evaluated either macroevolutionary or macroecological patterns of life history traits across several taxonomic groups, only few studies have attempted to integrate both dimensions in a single analytical framework. Here, we study how parity mode evolved across multiple scales in the radiation of spiny lizards (Squamata: Phrynosomatidae). We adopted macroecological and macroevolutionary approaches to explore how climate across spatial and temporal scales drives the evolution of viviparity in this lizard radiation. We find support for a weak signature of current climates on the geographical distribution of oviparous and viviparous species. By contrast, we detected that evolutionary transitions from oviparity to viviparity reach a peak during the Mid-Miocene Climatic Optimum — a period with a profound climate change event. We suggest that this abrupt climatic cooling promoted evolutionary transitions to viviparity simultaneously across three clades in the spiny lizard radiation. The decoupling in macroecological and macroevolutionary patterns found here suggests that past climate change has played a larger role than current climates in the spatial and temporal diversification of this reproductive life history trait.

## Background

The evolution of life history traits is one of the central topics in evolutionary biology [1–3]. Life history traits encompass a series of characteristics shaped by the interaction of extrinsic and intrinsic factors [2,4]. Several studies have examined how life history strategies evolved across the tree of life (e.g., mammalian life history strategies) [5–7] and how these traits respond to environmental variation [8–10]. Generalizations about these trait-environmental relationships have been reached in macroecological research [11,12], and macroevolutionary patterns of life history evolution have recently begun to be examined [13]. However, very few studies combine both macroecological and macroevolutionary approaches within a single framework, albeit some conceptualization about this integration has been achieved (e.g., [14,15]).

One of the most important life-history traits in squamate reptiles is the ability to produce fully formed live offspring [16–18]. Viviparity is considered an adaptation to survive in cold climates (“the cold-climate hypothesis”; [19]) and has evolved independently at least 100 times across Squamate reptiles [20–22]. Pyron & Burbrink [22] concluded that viviparity was the ancestral condition for all squamate reptiles, however this assertion has been strongly criticized from methodological and adaptationist perspectives [18,21,23,24]. Different lines of evidence support the hypothesis that viviparity evolved as a response to cold climates [19–21]. Thus, under the cold-climate hypothesis (hereafter CCH), viviparity is viewed as an adaptation to cope with low temperatures during the breeding season because gravid females, through behavior thermoregulation, avoid higher embryo mortality from cold temperatures [20,21,25,26]. However, there is ambiguous evidence supporting CCH [27–30]. For example, Andrews [27] and Shine et al. [30] found that nest temperatures of oviparous species were very similar to body temperatures in viviparous species, thus suggesting no differences in thermal regimes between these species groups. By contrast, Lambert & Wiens [29] provided support for a relationship between viviparity and low temperatures during breeding (egg-laying) season. Lambert & Wiens [29] also found that viviparity evolves more frequently in tropical mountain areas where temperatures are more stable throughout the year. These discrepancies warrants a further study exploring simultaneously how spatial and temporal dimensions, and its environmental correlates, have played a role in viviparity evolution.

It is well-known that temperatures in temperate regions vary strongly throughout the year, and that the breeding season in lizards usually occurs during the warmest season [25,31,32]. In contrast, tropical mountain areas exhibit temperatures consistently low year-round [31,32]. These differences in aseasonal and seasonal reproduction suggest a dichotomy between warm-cold temperatures that may not be a good proxy to test the CCH for clades encompassing a strong latitudinal gradient, such as spiny lizards. A multidimensional niche modeling approach might be more insightful to evaluate how environmental conditions drive viviparity evolution. For example, it is possible that marginal niche conditions—local niche conditions that are at the periphery of the available climate space—might be driving the evolutionary transition from oviparity to viviparity [33]. Marginal niche conditions might also be non-advantageous for offspring survival in oviparous species, and species might evolve to viviparity as an adaptation to cope with these non-optimal environments [34,35]. Including multiple climatic niche dimensions helps to understand whether species living at the periphery of the climate spaces tend to exhibit a given parity mode.

Phrynosomatid lizards are an excellent candidate system to explore how viviparity and diversification are linked. Some previous studies have tested whether aridity conditions have promoted species diversification in this clade [36]. For example, the number of co-occurring species tends to be higher in more arid areas [35], suggesting a strong effect of time in the speciation rates of phrynosomatid lizards, in which lineages have accumulated more in arid regions that have existed for longer amounts of time [35]. However, it is also possible that marginal niche conditions, given the climatic availability, have influenced the macroecological and macroevolutionary dynamics in this radiation of spiny lizards. Thus, the inclusion of more climatic niche axes (e.g., more variables) can help to discern how marginal environments drive current geographic patterns of life history traits [34,35,37].

Here, we combine macroecological and macroevolutionary approaches to examine how climate drives parity mode evolution in the radiation of spiny lizards (Phrynosomatidae family). First, we test whether cold climates have left a differential signature on species richness of viviparous and oviparous species. We expect to find a relationship between parity mode and present-day temperature, in which viviparous species are concentrated in colder regions (e.g., mountain regions) and/or more seasonal environments (e.g., temperate regions). In contrast, we expect oviparous species to be concentrated in warmer and more climatically stable areas (e.g., tropical lowlands). Second, we evaluate whether niche marginality and niche specialization based on a multidimensional niche concept (i.e., including several niche axes)[38,39] are correlated with parity mode. Accordingly, we expect to find viviparous species more concentrated toward marginal climatic niche space than oviparous species (Figure 1). Third, we examine rates of evolutionary transitions from oviparous to viviparous and vice-versa through time and how these transitions can be linked to climatic change events through the Cenozoic. We expect find higher transition rates coinciding with a known climate change events (e.g., Eocene-Oligocene Transition, Oligocene-Miocene Boundary, and the Middle Miocene Climate Optimum; [40]). Finally, we evaluate whether parity mode affects species diversification rates across the spiny lizard radiation. We expect to find that viviparity (as an emergent fitness trait; see [41]) promotes higher diversification rates in spiny lizards.

**Figure 1.**
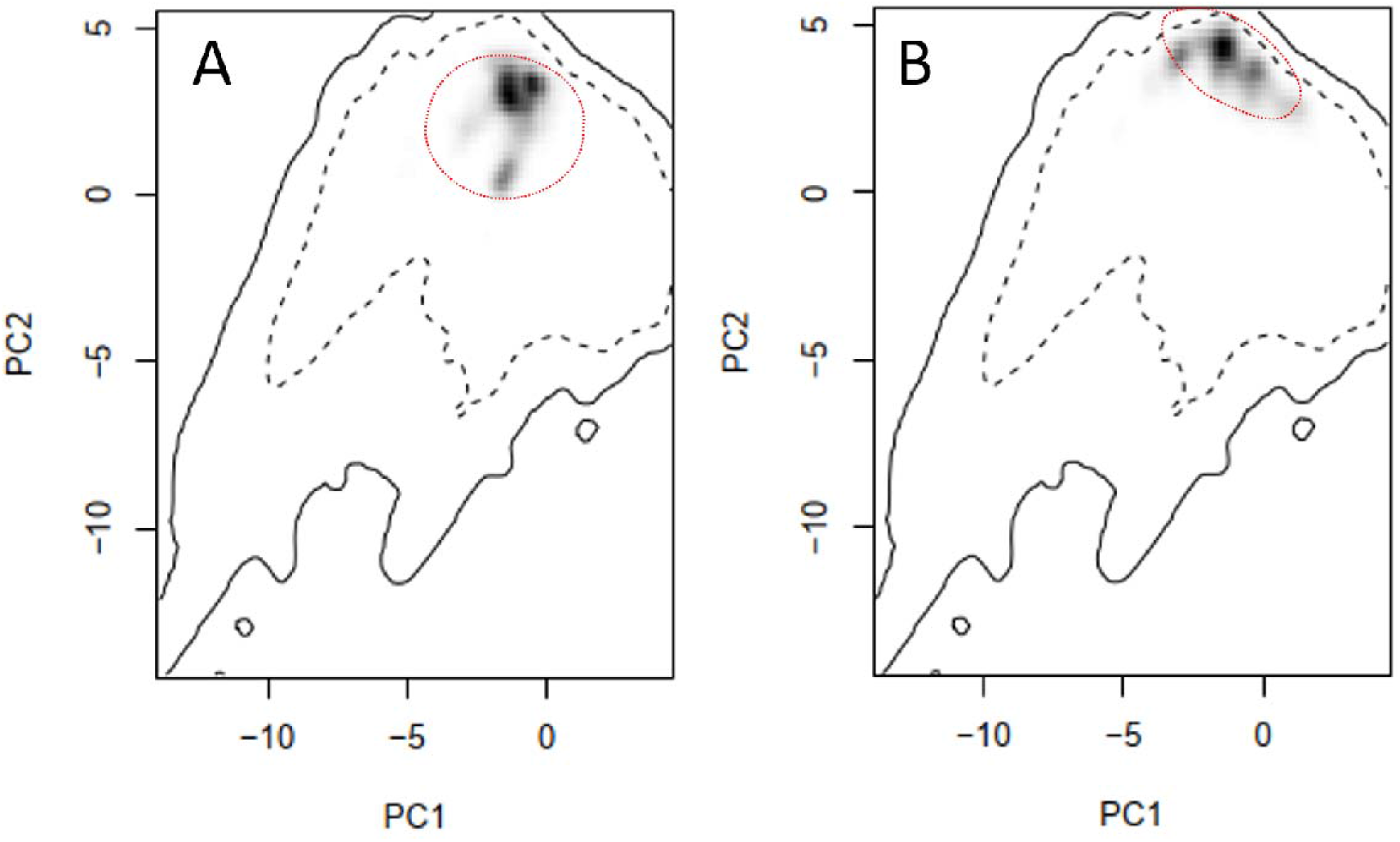
Hypothetical differences in species occupation patterns across climate spaces. In a) we represent species occurrences of a hypothetical species occupying central climatic niche conditions, delimited by dashed and continuous contour lines. In b) we represent species occurrences occupying marginal climatic niche conditions. We expect that viviparous species tend to occupy, in average, more marginal niche conditions than oviparous species.

## Methods

### Phylogeny

We used a time-calibrated phylogeny for the Phrynosomatidae family obtained from a large DNA dataset (i.e., 585 nuclear loci) from Leaché et al. [42]. This phylogeny includes 129 spiny lizard species (78% of the total known species; [43]) and it was built under a Bayesian approach and calibrated using several fossil records as calibrations points (see Leaché et al. [42] for further details).

### Life-history trait and occurrence data

We compiled information on the parity modes for all species in the Phrynosomatidae family using different literature sources (Table S1). We compiled occurrence records for all species (i.e., 164 species) from the Global Biodiversity Information Facility (GBIF) and cleaned up the records based on the known potential distribution of each species (e.g., [44–46]).

### Multilevel modeling approach

Using species’ geographical ranges [46] we generated presence-absence matrices (PAM) across grid cells of a regional grid with a resolution of 1^Q^ of latitudelongitude. This was done for oviparous and viviparous species separately. We mapped species richness across space for each parity mode by summing up the presence of species within a given grid cell. For each grid cell, we extracted two climatic variables from the WorldClim database: minimum temperature of the coldest month (bio6), and temperature seasonality (bio4) [47]. These two variables are considered the primary factors affecting geographical parity mode distribution according to the CCH. We generated a dataset with information from traits (i.e., parity mode, with oviparity coded as “0” and viviparity as “1”), climate (bio6 and bio4), and species presences (i.e., presence-absence of all spiny lizards) to test the “cold-climate” hypothesis. According to the “cold-climate” hypothesis, we expect that viviparous species richness increases toward colder and seasonal temperatures (i.e., in tropical mountains and temperate regions). By contrast, we expect to find that oviparous species richness increases toward warmer and stable areas (e.g., tropical lowlands). We used a multilevel modeling approach (MLM) using sites (i.e., cell sites) x species table and a site x environmental table (i.e., bio6 and bio4). This approach combines the assemblagebased and species-level approaches [48,49] in a single statistical framework, thus avoiding pseudoreplication effects and reducing type I errors (see [50] for further details). Statistical analyses were conducted using the *lme4* R package [51].

### Niche marginality and specialization

We generated ecological niche models for all species with more than five records using the ENFA algorithm [52,53]. This threshold of minimum occurrences is considered as the minimum number needed to fit relatively accurate ENFA models [52,53]. We fitted ENFA models with 19 bioclimatic variables from WorldClim. ENFA is a method that employs a factor analysis that make a comparison of the climatic niche of a species against the available background area. The first factor extracted is interpreted as a niche marginalization index, which is calculated as the multivariate ecological distance between the centroid of each species and centroid of the entire background area. The other factors extracted represent the specialization (i.e., the niche breadth), which is defined as the ratio of the variance of the mean background area and the mean for each species. We compared the marginalization and specialization niche metrics between oviparous and viviparous species. Finally, we tested whether parity mode was related to niche marginality and specialization using a phylogenetic logistic regression [54] using the *binaryPGLMM* function in the *rr2* R package [55].

### Ancestral state reconstruction and rate shifts through time

We inferred evolutionary transitions between states of parity mode through the spiny lizard phylogeny using several continuous-time Markov chain models (Mk models; [56]). We fitted four models as follows: 1) equal rates between states (ER model); 2) all rates different between states (ARD model); 3) irreversible model in which only transitions from oviparity to viviparity are allowed (IRRE1 model); and 4) irreversible model in which only transitions from viviparity to oviparity are allowed (IRRE2 model). These models were fitted using the *fitDiscrete* function in the *geiger* R package [57] and compared using AIC. We estimated ancestral states using 1000 histories of stochastic character mapping using the *make.simmap* function in the phytools R package [58] for the best-supported model. Finally, we visualized the rate of change in parity mode through time to evaluate whether rates were distributed homogeneously throughout the phrynosomatid lizard radiation or if they were concentrated in each temporal frame. We compared the observed pattern of parity mode evolution through time against a simulated scenario (null model) of both constant and equal rates between states through time, which was generated using the *ctt* and *sim.ctt* functions from phytools.

### Diversification analysis

We were also interested in testing the effect of viviparity on the diversification rates of spiny lizards, since it has been noted that viviparity provides evolutionary advantages for the colonization of cold conditions and consequently lineage diversification. We used a diversification model of the state-dependent speciation and extinction family (SSE), in which rates of speciation and extinction are related with the evolution of a binary trait—in our case we categorized viviparity = 0 and oviparity = 1—along a phylogeny [59]. To do so, we applied the Hidden State Speciation and Extinction model (HiSSE; [60]). This model incorporates an additional hidden character trait that allows the exploration of more complex scenarios, including unmeasured factors impacting diversification rates [60]. The HiSSE model evaluates the observed trait’s influence on the diversification rates of a lineage by estimating the following model parameters: net turnover (τ), extinction fraction (ε), rates of transition between the observed traits (q), and/or an unmeasured character trait. According to our species categorization (viviparity = 53 spp.; oviparity = 106 spp.), we fitted 24 diversification scenarios using a maximum likelihood (Table S2) approach, partitioned as: four under the classical Binary State Speciation and Extinction model (BiSSE; [59]), four under the Character-independent model (CID), and 16 under the HiSSE model with different constraints on parameters [60]. We selected the bestfitting scenario using Akaike information criterion under a pluralist approach by the joint estimation of ΔAIC, Akaike weights (wi) and evidence ratios (ER). This approach is a more concise way to quantify uncertainty in model selection [61,62].

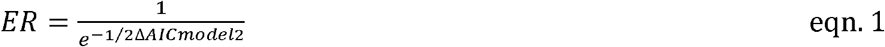

## Results

### Multilevel modeling approach

The patterns of species richness for all spiny lizard species did not exhibit a marked geographic pattern (Figure 2a). However, when we deconstructed the pattern according to parity mode, we find notable differences. Viviparous species exhibited a notable peak in species richness in areas that extend to the Trans-Volcanic belt in central Mexico (Figure 2b). By contrast, oviparous species exhibited at least three small peaks of species richness in temperate regions. The first one was located in the Sierra Madre Oriental in Coahuila; the second was located at the USA-Mexican border between Sonora and Chihuahua (Figure 2c); and finally, the third one was located in Baja California at the USA-Mexican border. The multilevel modeling approach reveals that only minimum temperature of the coldest month was significant, but only for viviparous species (Table 1). Finally, the species-cross approach revealed that viviparous species were concentrated in more stable climates (i.e., low temperature seasonality), but not in colder areas (Table 1]. These differences between approaches suggest that the signature of current climates on parity mode is not conclusive.

**Figure 2.**
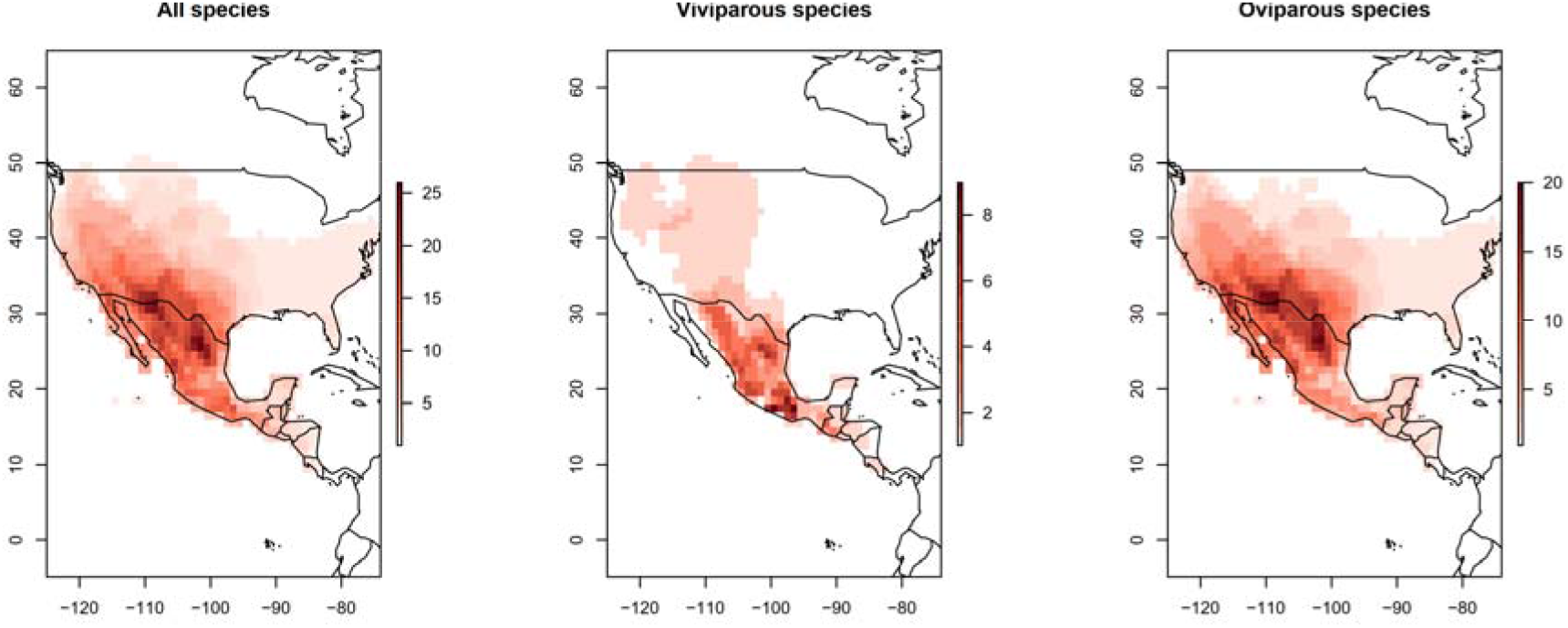
Geographical species richness gradients for phrynosomatid lizards, deconstructed for a) all species, b) viviparous species, and c) oviparous species.

**Table 1.** Model linking species presences, parity mode, and climatic variables across geography using a multilevel modeling approach (see main text for details). A trait-environmental relationship was tested while controlling for species and sites simultaneously. The only significant interaction was between minimum temperature of the coldest month (bio6) and parity mode. Marginal R^2^ values reveal a relatively weak climatic signature on trait geographic patterns.

| Fixed effects | z-value | p-value | Marginal R<sup>2</sup> |
| --- | --- | --- | --- |
| Intercept | -24.36 | < 0.001 | 0.122 |
| Minimum temperature | 9.878 | < 0.001 |  |
| Parity mode | 1.686 | 0.09 |  |
| Minimum temperature * Parity mode | -2.409 | 0.02 |  |

### Niche marginality and specialization

We did not find evidence for differences in niche marginality and niche specialization between viviparous and oviparous species (Figure S1; Table 2). Patterns of niche occurrence revealed that viviparous and oviparous species tend to occur in both central and marginal conditions of the total climate space (Figure S2). However, we discovered a potential association between marginal niches and viviparity in at least one clade of *Sceloporus* species (Figure S2). For the rest of viviparous species, we did not find an association between evolution toward marginal niches and a transition to viviparity mode.

**Table 2.** Phylogenetic logistic regression relating parity mode (i.e., a binary trait) against continuous niche traits (marginality and specialization) across the spiny lizard radiation. Climatic niche traits were calculated using 19 bioclimatic variables through an ENFA modeling framework (see main text for details).

| Climatic niche traits | z score | p-value | R<sup>2</sup> |
| --- | --- | --- | --- |
| Marginality | 0.5887 | 0.5560 | 0.021 |
| Specialization | 0.3956 | 0.3956 |  |

### Ancestral state reconstruction and rate shifts through time

The best-fit Mk model was one where only evolutionary transitions to viviparity from oviparity were allowed (Table 4). This model has relatively high support in comparison with an ‘equal rates’ model or an ‘all rates different’ model. Inferences of ancestral states with stochastic mapping reveal that the ancestral condition for all spiny lizards was oviparity, with only four independent transitions to viviparity (Figure 3). The first occurred in a small clade within the *Phrynosoma* genus. The second transition occurred in the branch descending to *Sceloporus bicanthalis,* a Mexican endemic lizard that occurs in high elevations (3000-4500 m) in the Trans-Volcanic belt. The third transition occurred in the clade forming the *Sceloporus formosus* group, a clade composed of both montane and lowland species occurring in tropical regions. Finally, the fourth transition occurred in a large clade composed of the *S. grammicus, S. megalepidurus, S. torquatus* and *S. poinsettii* groups. We did not find evidence of any evolutionary reversion to oviparity from viviparity (Figure 3). In addition, the highest mean number of changes between parity modes occurred during the Middle Miocene (Figure 4a). The highest ratio between mean number of changes and branch length also occurred during this time (Figure 4b). The observed rates of change show departure from a null model in which evolutionary transitions were fixed as homogeneous through time (Figure 4a, b). Accordingly, our inferences about evolutionary transitions being concentrated in a single short interval during the Middle Miocene are robust.

**Figure 3.**
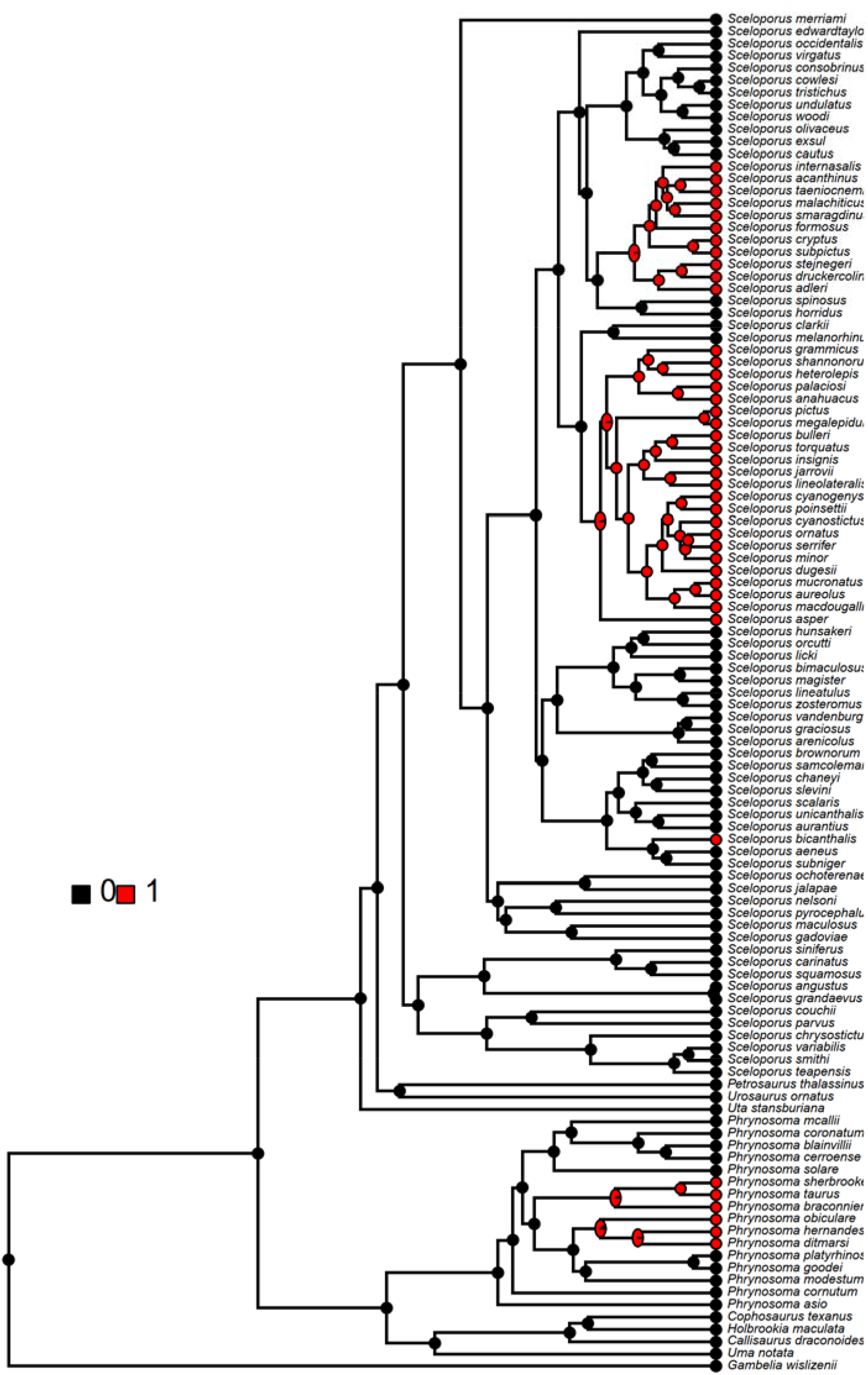
A summary of 100 histories of transitions to viviparity in phrynosomatid lizards using Bayesian stochastic mapping. Black: oviparous species (0); Red: viviparous species (1).

**Table 3.** Parameters of four Mk models fitted to parity mode data. Transition rates from oviparity to viviparity (q0->1), transition rates from viviparity to oviparity (q1->0), log-likelihoods (log(L)), number of fitted parameters (k), Akaike Information Criteria value (AIC). The best-supported model is highlighted in bold.

| Model | $q_{0 \rightarrow 1}$ | $q_{1 \rightarrow 0}$ | k | AIC | Akaike weights |
| --- | --- | --- | --- | --- | --- |
| All rates different | 0.004 | 0 | 2 | 53.71 | 0.20 |
| Equal rates | 0.004 | 0.004 | 1 | 53.34 | 0.24 |
| **Irreversible 0 → 1** | **0.004** | **0** | **1** | **51.63** | **0.56** |
| Irreversible 1 → 0 | 0 | 0.04 | 1 | 78.66 | 0.00 |

### Diversification analysis

Model selection suggests that there is not a single scenario in explaining diversification of spiny lizards (Table S2). Evidence ratios (ER) suggest that scenarios 8,18,15 and 10 are equally plausible for explaining diversification rates in this clade.

Among these four scenarios, scenario 8 (CID-4: ε’s equal, q’s equal) presents the lowest ΔAIC = 0 and the highest AICw = 0.1897 (Table S2), suggesting that diversification in spiny lizards is not linked to reproductive life history—either viviparity and/or oviparity—or that diversification is independent of this life history trait. The other equally plausible scenarios correspond to HiSSE scenarios, but these three scenarios assume that extinction fraction (ε) and transitions (q) are equal, like the character-independent scenario (CID-4). Thus, our results suggest that diversification of spiny lizards is mostly driven by hidden traits or that these hidden traits in conjunction with reproductive life history traits are influencing the evolutionary diversification in the phrynosomatid radiation.

## Discussion

We find contrasting evidence of geographic patterns of spiny lizard species richness according to a given parity mode. The species richness in viviparous species was concentrated towards tropical mountain areas, whereas oviparous species richness tends to be higher in temperate regions. It has been widely supported that viviparity evolved as a response to low temperatures according to the cold-climate hypothesis (CCH). This hypothesis has gained support by the fact that viviparous species tend to occur in tropical mountains where temperatures are stable and low all year round. Although we find an association between viviparity occurrences across geography and colder temperatures, the marginal R^2^ values were relatively low, suggesting a weak geographic effect. Furthermore, we did not find evidence that viviparous species tend to occupy marginal niche conditions or have narrow climatic niches. Accordingly, the inclusion of more niche axes (i.e., more climatic variables) has affected the inferences on trait-climate relationships. By contrast, we find that evolutionary transitions from oviparity to viviparity reach a peak during the Middle Miocene Climatic Optimum (MMCO) period (Figure 4). This period, which occurred ~14 mya, is well-known as an abrupt climate transition towards colder temperatures [63]. It is possible that a rapid cooling during this time acted as a trigger of viviparity diversification, and the resulting geographic pattern of viviparous species (i.e., with more species toward colder, but not seasonal, climates) is a direct response of that evolutionary legacy rather than present-day conditions.

**Figure 4.**
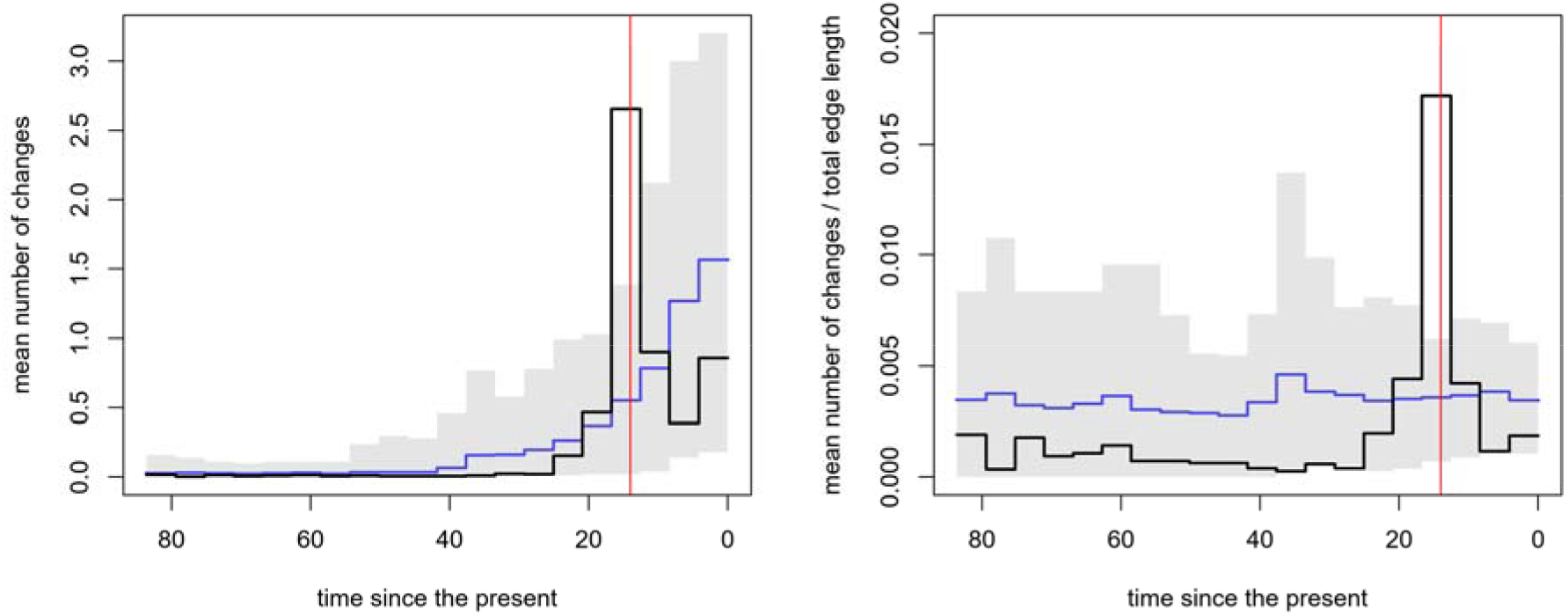
Changes in parity modes through time in Phrynosomatidae lizards, a) Mean number of changes inferred through time, b) Ratio of changes through time. Vertical red line represents the Middle Miocene Climatic Optimum event (Young et al. 2009).

The concentration of shifts toward viviparity in a small temporal frame, characterized by an abrupt climate change, provides support for the link between cooling temperatures and viviparity emergence. It remains a possibility that each of the four transitions toward viviparity as inferred in our study was driven by different factors (e.g., colonization of mountain areas and/or shifts in reproductive cycles among species). As the four evolutionary transitions explain the entirety of viviparity diversity observed in the phrynosomatid radiation (i.e., 53 of 156 species), it might be necessary to evaluate whether similar physiological and developmental mechanisms are underlying each transition across the phrynosomatid clade. For instance, it would be useful to establish whether similar egg retention times and earlier initiation of reproduction can be identified as exaptations [64] to viviparity in the four independent transitions inferred in this study.

Viviparity evolution in *Sceloporus* implies at least three adaptative shifts in evolutionary regimes [32]. Méndez-de la Cruz et al. [32] suggested that the first regime shift during the transition from oviparity to viviparity was associated with colonization of mountainous cold regions. The second regime shift corresponds to a major shift in the timing of the offspring birth from autumn to spring. Finally, the third one involves a regime shift in the reproduction cycle from asynchrony to synchrony in male and female mating. We were able to infer three independent origins of viviparity through the spiny lizard phylogeny, but to test the Méndez-de la Cruz et al’s hypothesis it i=s necessary that the other two trait transitions (i.e., shift in breeding time and mating and breeding synchrony) occurs simultaneously. The synchrony between parity mode, breeding time, and mating synchrony across the spiny lizard radiation needs to tested under a phylogenetic comparative approach (see [62] for an example].

We suggest that macroevolutionary patterns of reproductive life history traits in spiny lizards are decoupled from current environmental conditions. Lambert & Wiens [29] found evidence of current climatic effects on evolutionary patterns of viviparity in spiny lizards. We found evidence for a single variable (i.e., minimum temperature) explaining the trait-climate relationship. When more niche dimensions were included and marginal niche conditions were explored for viviparous species occurrences, we found no such association. The concentration of viviparity transitions in a relatively short time (~14 mya; Figure 5) suggest that past climatic change events have likely played a greater role in trait evolution in spiny lizards than current climates. It would be interesting to explore whether this association between past climatic changes through Cenozoic [65,66] occurs across the entirety of Squamata. In addition, the lack of a relationship between parity mode and species diversification suggests that this life history trait is not leaving an emergent fitness effect [41] in spiny lizard species. Finally, by the simultaneous evaluation of spatiotemporal dimensions of life history traits under macroecological and macroevolutionary approaches, it is possible to disentangle the role of current and past climate in shaping life history traits and their macroevolutionary effects.

## Data accessibility

Data and R codes are available from the Dryad Digital Repository:

## Authors’ contributions

J.A.V conceived the ideas, analyzed data, and lead the manuscript writing; G.C.-G compiled life history traits, provided ideas, and checked taxonomy for species datasets; J.P.-L analyzed data, provided ideas, and drafted the manuscript; O.F.-V provided ideas and drafted the manuscript. All authors revised and approved the manuscript. All authors agree to be held accountable for the content of this paper.

## Acknowledgements

We thank Brett O. Butler for revising the English. We would like to thank the anonymous reviewers and the editor for helpful comments.

